# Practice of Cardiac Auscultation: Clinical perspectives and its implications on computer aided diagnosis

**DOI:** 10.1101/013334

**Authors:** Subhamoy Mandal, Roshan J Martis, K M Mandana, U Rajendra Acharya, Jyotirmoy Chatterjee, Manjunatha Mahadevappa, Ajoy K Ray

## Abstract

Study of the disease demographics in human population indicates that cardiac ailments are the primary cause of premature death, and a need for emergent technologies is felt to address the rising trend. However, development of automated heart sound analysis system and its usage at the grassroot levels for cardiac pre-screening have been hindered by the lack proper understanding of the intrinsic characteristics of cardiac auscultation. In this article we present an investigatory report based on a nationwide survey conducted on the practice of cardiac auscultation for determination of its effectiveness in diagnosis. The aims are to achieve better validation of heart sound acquisition methods and use the clinical feedback from cardiologists for improvements in the classification of the cardiac abnormalities. Results obtained from six different classifiers used in the study are illustrated, which show a remarkable specificity using an improvised classification hierarchy, derived based on clinical recommendations. The study addresses the needs for better understanding of the relevancy of heart sound signal parameters, recording transducers, recording location and the inherent complexity associated in interpretation of heart sounds, specially in noisy environments of out-patient departments and primary healthcare centers. Further, the inter-relationship between heart sound and other advance medical imaging modalities, and the need for more focused training in cardiac auscultation among the medical and paramedical staff is investigated.

## I. INTRODUCTION

AUSCULTATION provides important clinical information and cost effective measures to determine treatment strategy. Conventional wisdom mandates the use of auscultation as a compulsory tool for physical examination and selection of additional tests that will provide clinical information of independent value, at no significant cost. Auscultation is practiced for evaluating the functional status of a complex human system such as the heart, lungs, abdomen etc. In the current study, our particular interest is the auscultation of heart, generally referred to as *Cardiac Auscultation* or *Heart Sounds* (HS)in medical literature [1]. The HS are reflective of hemodynamical processes of the heart, are often barely audible, and involves dominant frequencies at the lower frequency edge of the threshold of hearing. They are brief, transient sounds, and it is hard to interpret the exact sequence of cardiac events and to isolate them. Cardiac auscultation is necessarily subjective, and the inherent difficulty in hearing and correctly interpreting heart sounds offers suitable opportunity to use signal processing and machine learning systems to aid clinical proceedings. The auscultatory sounds of hearts comes multiplexed with the lungs sound and additional movement artifacts, the filtering and denoising steps incorporated in the system designed are thus vital for segregating the desired signal and enabling automated cardiac prescreening [2].

Auscultation has been used by physicians as a valuable clinical tool for cardiovascular evaluation. The general belief in the medical community is consensual about the applicability of HS analysis to screening and treatment planning, however, studies indicate towards the declining practice and teaching of these techniques at all level of training [3]. This is attributed to the emergence of newer modalities which provide the same information, thus undermining the importance of HS evaluation [4]. However, automated analysis of HS, especially towards the design of effective screening mechanisms, have been generally neglected. Most research articles have, nevertheless, demonstrated the vitality of the ausculatory methods. Mangione et. al. [4] stressed that the over-dependence on modern technology is attributed to lack of training and motivation. The trend further amounts to loss of valued medical knowhow and economically strains the one paying for the health services. A suitable device designed to optimize the information gained through auscultation, and overcome the shortcoming of existing tools is a need of the hour. The device can be of significant value in screening and advanced test selection in cardiac conditions, specially in economies having low per-capita healthcare expenditure [5], [6].

The current study aims to establish the effectiveness of HS in rural setups lacking advanced imaging and Doppler facilities, and evolve a standardized framework for acquisition and processing of HS signals. Moreover, archival recording of heart sounds is not currently part of standard clinical practice, and no standard instrumented auscultatory investigation is followed to support referrals to echocardiography. We have tried to accumulate feedback from medical community and design a suitable framework for promoting computer assisted cardiac auscultation [7]. In this article we have briefly summarized the science behind auscultation and included the results obtained from a nationwide survey which throws up new insights for suitable addressing the defined goals.A new scheme for classification of HS is also illustrated, the scheme aids in translating clinical acumen into computer-aided prescreening methods.

## II. Clinical Significance and State of the Art

Auscultation represents the acquisition of mechanical vibrations from the surface of the body that encompass the frequency range of sound; vibrations below this range (20 Hertz) are palpable and constitute a source of supplementary information (*as defined by* [3], [8]).

*Practice of Clinical Cardiac Auscultation:* Auscultation is practised for studying the cardiovascular, pulmonary and gastrointestinal systems. The focus of the study is strictly limited to cardiovascular system, and the heart to be specific. PCG represents the modern form of cardiac auscultation based on graphic recording of the HS and the murmurs. Accurate timing information is maintained by simultaneous carotid pulse recordings giving the onset of aortic ejection and valve closure and also by simultaneous ECG showing movements of valves without phase shift [9]. Auscultation is an integral part of internal medicine training and widely discussed in medical texts. [10] outlines the basic physiology behind the genesis of HS and its’ relationships with ECG and other cardiac parameters. The clinical aspects of auscultation have been illustrated by [1], it also describes in details the relation of HS with corresponding ECG signal. [9] is a comprehensive text rendering the basics of HS and the indicators for its’ identification; it was extremely helpful in the basic understanding of the methods. [11] attempted to address the question whether a Doppler Echocardiogram is needed to be used routinely or whether auscultation is sufficient for diagnosis of functional murmurs. We also referenced the work of [4]; the work surveyed U.S. internal medicine programs to assess the time and importance given to auscultation training, and the auscultatory proficiency of medical students. Tavel et. al. [8] explores the perspective of intervention of signal processing and biomedical devices to overcome the deficiencies in traditional auscultation. It argues that HS analysis is a cost effective means of additional treatment planning, and provides impetus for technical advances to address the challenges.

*Technological Developments:* In literature we find references of signal processing based approaches for HS analysis since 1988, and thereafter, there have been sustained contributions to this domain. Electronic stethoscopes have been available for quite a long time, but they offered marginal improvements over the mechanical predecessors. They were not very successful because of its high cost without much additional benefits and, susceptibility to noise from external sources. Moreover, most models lacked the facility to store and later recall the sound for effective differential diagnosis, and conveniently transmit the sounds to remote sites. With the development of processor and communication technologies this gap has slowly been bridged [3]. The electronic stethoscopes currently available in the market (HD Medical, HDfono, 3M Stethos) fulfil the basic criteria in terms of signal quality and connectivity, but are designed for use by trained physicians only or for educational and training purposes. However, a screening device based on these methods is conspicuously unavailable, and research efforts towards achieving a framework for cardiac prescreening and diagnosis using HS are rare. CadioScan (Zargis) is the only example of FDA approved model that assists primary care physicians in the interpretation of HS and referral of patients with heart murmurs [5]. Reed et. al. [12], [13] proposed a model based approach which finds prominence in conceptualization of system depicted in the dissertation. [12], [13] designed a model for HS generation and characterizing them based on the model for determining clinically important feature. They also presented a prototype system for HS analysis using wavelets and neural networks based classifiers. Syed et. al provided an insight into the relative difficulty of various steps involved in accurate interpretation of acoustic cardiac signals [14] and compared the efficiency of different workflows. The current project aims to design a system which can be conveniently used by paramedical staff (including auxiliary nurses, midwives and traditional ‘bare-foot’ doctors) for screening of cardiac cases in absence of trained cardiologists. A more detailed study of the hardware system and the signal processing methods developed are illustrated in Mandal et. al. [2]. The new system incorporates a denoising blockset and novel customizations for comprehensive processing of cardiac signals in noisy outpatient conditions, an improvement on earlier frameworks that allows the system to be used in rural health centers as well as mass screening operations [15].

## III. A Nationwide Survey on Practice of Cardiac Auscultation

In India, a comprehensive survey on cardiology screening has not been conducted extensively till date. A distinguished pediatric cardiologist and our panelist, stressed on the fact that currently our statistics are based on surveys conducted in developed countries, thus the significance of an indigenous survey is immense. This led to the initiation and execution of the questionnaire based survey entitled, “*A Survey on Practice of Cardiac Auscultation for Determination of Its Effectiveness in Diagnosis*.”

### A. Survey Design

The study designed and conducted is a unique attempt to identify the efficacy, teaching pattern and relevance of auscultation in cardiac pre-diagnosis. The main purposes of the survey are identified as below:

- Identify the general practice and variations in methodologies for auscultation.
- Characterizing the timing and diagnostic features of Heart Sounds in clinical practice and its applicability in computer aided analysis.
- Ascertain the level of training and expertise in auscultation during medical education and its effect in diagnostic decision making.
- Study the inter-relation and relative efficiency of ECG, PCG, Treadmill test (TMT) and Real-time Echocardiography (RT ECHO).

The information collected through this questionnaire based survey aims to identify the variables in the practice and teaching of cardiac auscultation. It helps us to recognize the diagnostic significance of the cardiac auscultation and the methodology to be adopted; and to establish its suitability for computer aided analysis. We will also try to comment on efficacy of multiple diagnostic modalities in unity or unison through the responses.

The survey comprises of seven (7) independent sections, the sections first through fifth are complimentary to the defined goals are in question-response format, special comments and respondent's details are accommodated in the sections 6 and 7 respectively. The first section asks the respondents to specify the diagnostic tools used and the related experience, personal confidence level with the modalities, teaching methodology in cardiac auscultation training adopted and the current access to the monitoring modalities. The second section addresses the query on the timing information (systole/Diastole-Early, Mid, Late), positions of auscultation, sensor used, pitch (frequency) of sound, and special focus in interpretation of diastolic murmurs. Third Section asks the opinion of the respondents on the importance of HS of in clinical diagnosis and whether more time needs to be devoted to during cardiology training. The fifth section asked for rating the importance of identification eleven cardiac (abnormal) events. The 11 cardiac event selected were:

1. S3 Gallop
2. S4 Gallop
3. Early systolic Click of aortic ejection
4. Mid systolic Click of mitral Valve prolapse.
5. Opening snap of mitral stenosis.
6. Pericardial friction rub
7. Systolic murmur of mitral regurgitation.
8. Systolic murmur of aortic stenosis
9. Diastolic rumble of mitral stenosis.
10. Diastolic murmur of aortic regurgitation.
11. Continuous murmur of patent ductus arteriosus

For each event, the participants had to select one of the five predefined linguistic significance levels - i) Strongly Agree (4), ii) Somewhat Agree (3), iii) Neutral (2), and iv) Disagree (0).

The fourth and the fifth sections were exploratory in nature, aiming to address the inter-relationship of modalities, their applicability, and the importance of technological interventions in providing better diagnostic support. Earlier survey conducted by [4] on practice of cardiac auscultation in internal medicine and cardiology teaching for accredited U.S programs was followed while drawing up the questionnaire and suitably improvised to address the Indian medical fraternity. We also followed the “Guidelines on Assessment and Management of Cardiovascular Risk for Medical Officers” developed under the Government of India - WHO Collaborative Program [2008-2009].

*Target Experts:* The provision of the information was voluntary. The questionnaire was accompanied with a detailed statement of purpose (SOP) to appraise the participants and encourage them to provide accurate information. Consistency between the practice and responses were needed to make the survey results more representative and substantiate its use as the basis for statistical evaluations. Thus, we made the respondents aware of our methodologies and system design effort through the Statement of Purpose and sharing key manuscript(s). In many cases we interacted directly with them over phone or email explaining further details about the survey requirements, and clarify the doubts arising during filling in the responses. We had invited eminent cardiologists, cardiothoracic surgeons, medical practitioners and residents from across the country. A total number of fifty questionnaire packages were sent to selected individuals with a letter of invitation; we recorded responses from thirty two individuals. The confidentiality of the information respondents provide is pledged to be carefully protected and only aggregate information and no details of individual responses will be released as part of the survey results. The respondents to this survey had a the right of access and correction to their submitted responses for a short period (12 weeks), post expiration of the period or voluntary opt-out the had access to the consolidated results of this survey.

### B. Report on the Survey Outcome

For the current article, we have scored twenty-one responses based on two categories; nine responses were from undergraduate students and were not considered for the study, and two responses were not considered due to incomplete data and inconsistencies. The survey respondents were classified under two categories (A) Experienced Cardiologists having experience 6 years and structured training in cardiology; and (B) Trainee Cardiologists having experiences 6 years and might or might not have undertaken a comprehensive cardiology program. We recorded a participation of professionals from 13 states/Union Territory of India (Andhra Pradesh, Chandigarh, Karnataka, Kerala, Madhya Pradesh, Maharashtra, Punjab, Delhi, Orissa, Tamilnadu, Rajasthan, Gujarat and West Bengal). Individual reporting experiences of the participants are in the range of four to thirty five years with average reporting experience of 12.4 years for category A and 3.7 years for category B.

The Auscultation score was computed using the linguistic scores assigned by the doctors and scaling the same with the experience levels (see III-A). Mathematically, it can be expressed as,

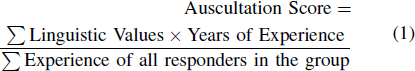

To ease the process of sending the responses both online and postal methods of returning the evaluation sheets were made available to physicians. A format of the Questionnaire is available in as supplementary material.

*General Information:* The section indicated that the physicians primarily use auscultation and ECG as prescreening tools, with 92% of the respondents agreeing to it. RT ECHO was the next with 66% consensus. Computed Tomography (CT) was the least used modality with score less than 40%. Most of the participants (84%) have undergone structure cardiology training during their undergraduate of specialization studies, the preferred method of imparting the education being through Lectures and demonstration. Audio cassettes and recordings were also acknowledged as helpful tools.

*Auscultation Practice:* The participants commented on the timing of the HS and we got a balanced response for the entire cardiac cycle (both systole and diastole) with focus of listening early stages of same, the importance decreases gradually with mid and late systolic and diastolic phase [Refer Figure 1].

**Fig. 1.**
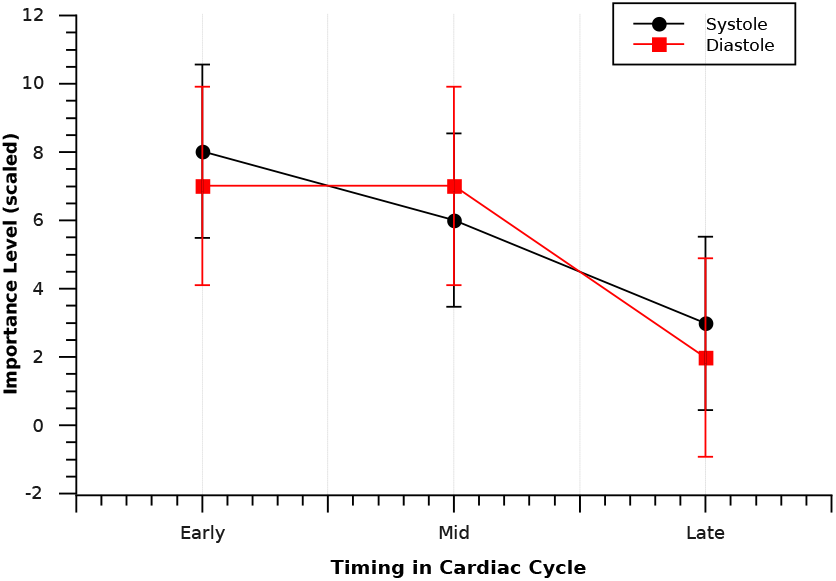
Importance of Timing Information of HS Components

The second important question posed was regarding the positions of auscultation. This is much debated and threw up many different suggestion and observations. After carefully studying the input we were able to finalize the following standardized location to be used for acquisition of HS:

- 4th ICS Mid axillary line left side - levocardia
- 5th intercostal space midaxillary line
- 2nd intercostal space left sternal border
- Murmurs can be detected in infraclavicular region and back.

The ECG locations are not found to be useful for auscultations, but the areas corresponding to mitral, bicuspid, tricuspid, pulmonary and aortic regions are to be monitored; murmurs can be heard in all these and even beyond.

The third important issue that the respondents commented on was the pitch (frequency) of the HS they concentrate on. The four frequency levels viz. very high, high, moderate and low were given equal stress with focus being more on diastole by 35% in all observations. The construction of the stethoscope plays a vital role this context. The chest piece of the stethoscope have a rigid diaphragm and a bell. The diaphragm allows better audition of the high frequency sounds, e.g. splitting sounds, opening snap, aortic diastolic murmurs. The bell transmits the low frequency sounds better, when applied lightly to chest, e.g. ventricular filling sounds and mitral diastolic murmur [9]. The responses show, diaphragm was the most used mode for picking up the sounds (78%) closely followed by Bell (68%). Majority of experienced participants indicated the use of Bell for picking up of low pitch sounds and diaphragm for high pitch sound sequentially in a single investigation. This finding is important in design of the transducer module, but is currently out of scope of the thesis.

*Auscultation Training:* Two of the more important questions of the exercise inquired the participants were: (1) Importance of auscultation as a clinical tool in the current scenario, and (2) more time should be devoted to teaching of auscultation during internal medicine/cardiology training.

It is observed that the experienced cardiologists give more weightage to ausculatatory skill compared to newly trained doctors (see Figure 2). The experienced often place HS as one of the primary tools for screening and diagnosis, whereas the trainees seem to regard it more as a mandatory secondary tool. The trainee doctors felt that the knowledge imparted through classroom teaching and demonstrations are a necessary component of the internal medicine practice, there is a slight bias towards more training. The experienced group, however, believe that the present training is at-large insufficient and the trade can only be mastered through sustained Out-patient Department (OPD) practice.

**Fig. 2.**
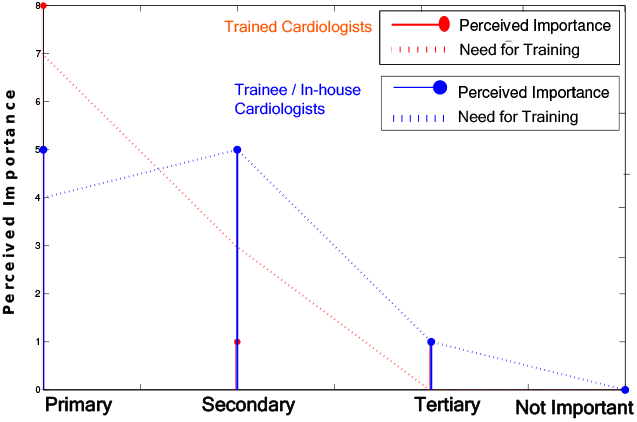
Importance of Heart Sound Components based on Timing Information

In the fifth section, we had asked for the feedback of the respondents on the 11 different cardiac events as mentioned in the survey design. The ranked responses using weighted sum (linguistic level) for the events were scored for both the categories. The results of the same can be depicted by the Figure 3.

**Fig. 3.**
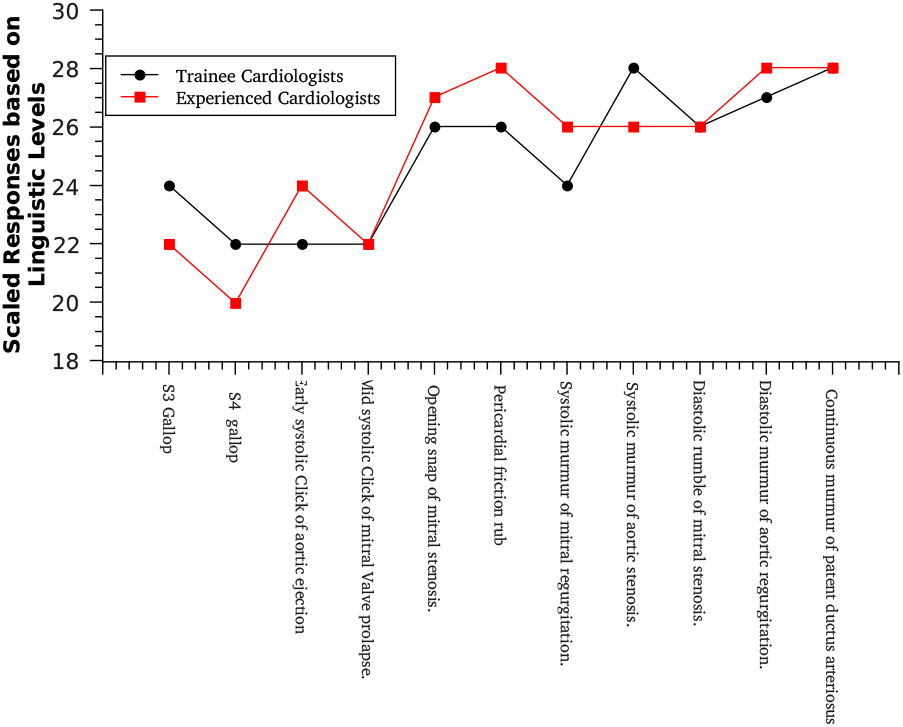
Perceived importance of different HS in auscultation training

We conducted the Anova test based on the weighted responses to the 11 different cardiac events. The test reveals that the median for the experienced observers is relatively higher than that of the trainees. Moreover, the responses have been spread over a greater range (high intra-class variance) for the trainees. This clearly indicates that the experienced cardiologists are more focused while using HS as a tool and confident about choosing the useful features (Figure 4). This quantifies the usefulness of experience and basic clinical acumen in HS analysis.

**Fig. 4.**
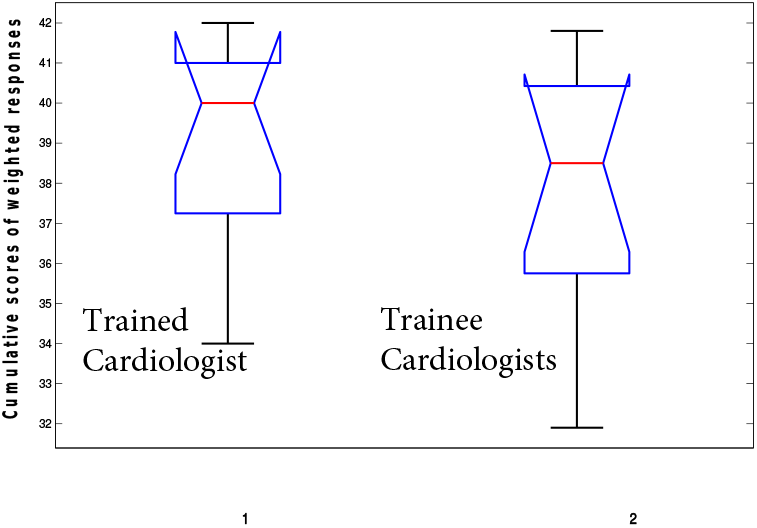
Cumulative scores distribution of responses by (1) Experienced cardiologists and (2) Trainee Cardiologists for determining importance of 11 selected Cardiac events

*Multimodal Applications and Future Directions:* Due to the structural similarity between the questions and the responses in the fourth and fifth sections, we have consolidated the results together. Section five inquires if use of multiple modalities will greatly influence cardiac screening; 90% of the experiences and 82% the trainee cardiologists have agreed to the same. The respondents were also asked to rank the combination of modalities based on their efficacy, the combination of auscultation and RT ECHO was the most preferred option for detecting murmurs, and combination of TMT, auscultation and RT ECHO was perceived to be most useful for detection of coronary artery disorders. It is to be noticed here that, though the contribution of ECG is well appreciated in recent research [16], it was not reflected in the physicians responses. ECG performed rather poorly in both the cases, and cardiac auscultation was found to be very promising with second and fourth ranks for the purposes mentioned. It is, however, agreed that ECG finds importance in management of post-stoke subjects. It also forms an component of Doppler echocardiography and Tissue Doppler Imaging for detailed investigations, diagnosis and cardiac rehabilitation. The cardiologists were asked if the use of better equipments can enhance the performance of diagnosis using auscultation, the option seemed to be very undecided and debated. While 40% believed it will strongly influence performance, 40% were neutral to it, 10% said it will marginally increase the performance, the last 10% believed that it will have no or even adverse effects on the skill. Finally, only 20% reported to have found HS important for coronary artery disorders. This was an important reporting since modern input coming from biomedical engineering domain proves the usefulness of HS in the identification of CAD using newer sensors and picking up data well beyond the lower threshold of human audition [17].

### C. Comparison of the findings

The results have been found to be consistent with the investigations done by [11], which opines that well trained cardiologist can identify the vast majority of murmurs on auscultation. This can prevent the over usage of Echocardigraphy and reduce medical expenses. The key markers for the survey was compared with a similar survey conducted by [4]. The general trend in both the survey were found to be similar, though variations exists. It is heartening to see that the perceived importance of HS by both peer groups in India and U.S. is rated as high on average. It is also encouraging to note that in India, 84% of doctors receive some sort of structured training pertaining to auscultation. In comparison, only 37% of U.S. internal medicine trainees receive a structured training (76% for cardiologists). The perceived importance of different HS in training (Figure 3) by both trainee and experienced groups have been found to be coherent, such cannot be established for the results given by [4]. The surveys presented divergent trends when continuing medical education was taken into consideration. [4] reported 74% cardiac fellows retraining on auscultation, whereas in India, the trend is more prominent towards enhanced usage of sophisticated modalities (72%). Even though its a growth marker, it amounts to significant loss of acumen in usage of traditional methods. The survey significantly contributes towards evolving a scheme for classification of signals based on medical feedback. It also helps in understanding the cardiac healthcare needs and its variation in the Indian subcontinent. This study being the first of its kind in the geographical area, and its ethnography being unique from the earlier surveys, it deserves special attention.

## IV. Classification of Heart Sounds

Based on the medical feedback we came up with a modified classification scheme which accounts for frequency properties of the cardiac abnormalities and mimics the action of a physician. The system initially distinguishes between (a) **Normal** and (b) **Abnormal**, and then sub-classify abnormal signal samples into physiological and anti-physiological (or pathophysiological) flow conditions. For detection of the normal signals the system checks for integral S1 and S2 sound (not split), absolute frequency of sound varies between 40-250 Hz. The murmur sounds are generated by turbulent flow of blood, thus the nature of sound directly depends on the turbulent flow properties. This alteration of sound signature helps us detect the abnormal conditions (using frequency analysis techniques). We observe that the abnormal flows are primarily of two types-(i) Physiological flow that is typically seen in stenosis, the blood flow is turbulent but it still flows the physiological direction. (ii) Patho-physiological: The blood flow is turbulent e.g regurgitation and part of the blood flows in direction opposite to the natural directions. These changes cause occurrences of very specific distinguishing frequency properties, the frequency profile for regurgitant murmurs being sharper and shorted. The Figure clearly indicates the difference in profiles of the stenosis and regurgitation. There are certain disorders not to be identified by this method; we classify them as a separate abnormal-undefined class. This abnormal undefined class is prone to higher chances of error; however, this scheme performs with greater accuracy when employed to screening problem. The method also enables us to go for a structured method to delineate to abnormalities and employ advanced signal processing measures to give more accurate interpretation. The frequency and temporal cut-off for the classification was fixed after thorough investigation of frequency profile of HS signals in the entire databases and wide consultation with collaborating doctors.

### A. Classification Methods

The typical normal ECG with QRS middle point and the corresponding PCG signal are depicted in Figure 5. The PCG signals are subjected to dimensionality reduction using Independent Component Analysis (ICA) [18] [19] [20] [21] [22] [23] [24] [25] followed by pattern classification. In this study, Support Vector Machine (SVM) [26] [27] with different kernel functions (quadratic, polynomial and Radial Basis Function (RBF)), Decision Tree (DT) [18] [19] and Probabilistic Neural Network (PNN) [18] [19] are used as classifiers. The tenfold cross validation is used to calculate the performance of classification.

**Fig. 5.**
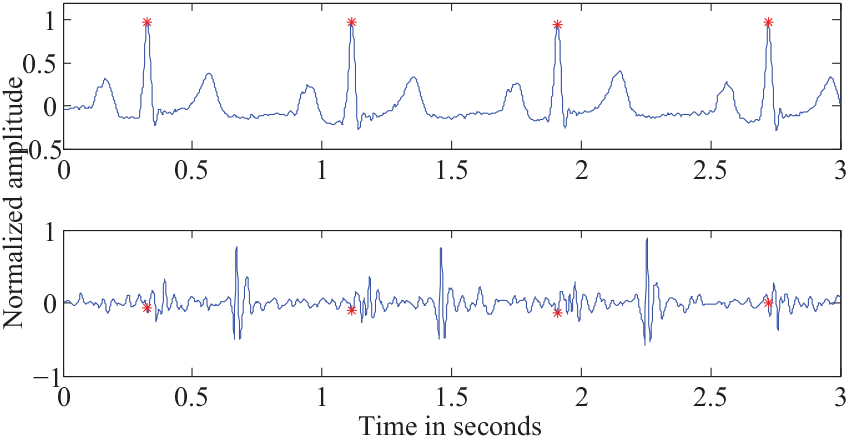
Normal ECG and PCG signals with QRS complex middle point shown in red asterisk

### B. Results of Classification

The PCG signals from normal, Aortic Stenosis (AS) and Aortic Regurgitation (AR) classes are used in this analysis. In total 145 normal, 102 AS and 52 AR segments of PCG are used. The noise and unwanted artefacts are removed using appropriate filtering techniques. Since there are a large number of samples in each signal, the dimensionality reduction using ICA is applied. After ICA ten independent components are chosen and are used for subsequent pattern identification. The classification is performed using ten fold cross validation. The average performance (sensitivity, specificity, PPV and accuracy) is shown in Table 2 for different classifiers. It can be noted from Table 2 that all classifiers provide 100% average specificity. It implies that in each of the ten folds, all normal patterns are classified into normal. Hence this proposed system can identify the normal patterns correctly in all cases. If it is not so, the patient who undergoes screening with this developed technology will unnecessarily undergo mental trauma. As seen from Table 2, the DT provides 82.6% of average accuracy, sensitivity and specificty of 100% and 98.57% respectively.

**TABLE I.** Responses to the Questionnaire on Cardiac Auscultation

| Evaluator | Response Categories |  |
| --- | --- | --- |
|  | Cat A | Cat B |
| Total Responses | 9 | 12 |
| Avg. Experience | 12.4 | 3.7 |
| Auscultation Score | 3.71 | 3.34 |
| Preferred Modality | PCG+ECHO | PCG+ECHO+ECG |

**TABLE II.**
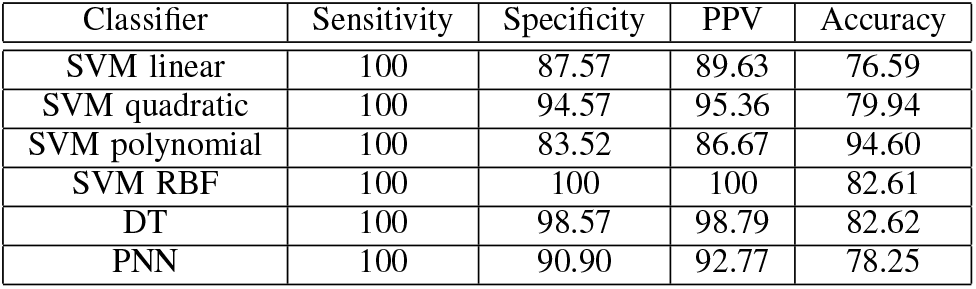
Classification Performance

| Classifier | Sensitivity | Specificity | PPV | Accuracy |
| --- | --- | --- | --- | --- |
| SVM linear | 100 | 87.57 | 89.63 | 76.59 |
| SVM quadratic | 100 | 94.57 | 95.36 | 79.94 |
| SVM polynomial | 100 | 83.52 | 86.67 | 94.60 |
| SVM RBF | 100 | 100 | 100 | 82.61 |
| DT | 100 | 98.57 | 98.79 | 82.62 |
| PNN | 100 | 90.90 | 92.77 | 78.25 |

## V. Conclusion

Cardiac auscultation is a part of general case taking, and thus all physicians practice it, but in India the use of HS for organic disease diagnosis is limited to experienced cardiologists only. There has been an increasing trend of usage of advanced modalities like Doppler Echocardiography, CT and MRI are employed without proper screening. Auscultation can act as sufficiently sensitive tool for identification of systemic murmurs disorder and screening of subjects [28]. In case of prescreening and health visits (including chamber practice) a good practice of HS evaluation couple with patient history can be indicative of the underlying pathogenesis. There has been a waning interest in traditional understandings of HS owing to over reliance on advance technologies. This in turn is a contributory factor to ever increasing healthcare divide with most physicians preferring to stay on at urban locations setting lack of facilities [2]. Re-inculcation of the traditional diagnosis skill and suitable training of paramedical stuff and offering them with innovative devices (as one proposed in [2]) is vital to strengthen the health system [5]. This article consolidates the results of the cardiology survey, the suggestions acquired by interviewing eminent cardiologists, and the knowledge gained through sustained interaction with the physicians. This new information was skillfully exploited in designing new classification schemes which shows remarkable specificity. Further, we were also able to show how the interplay of different modalities can positively reinforce the results.

In a scenario involving developing countries like India, it is to be understood that the advanced modalities are available only in the tertiary care centres. As one of our participants aptly points out, “The primary care giver is bereft of any such facility, and will have to do with clinical ‘eye’ and skill of ECG and auscultation. Importantly, they are the primary screener - tertiary care givers are mostly implementers of definitive managements.” The primary health system should, thus, be made sufficiently strong so that morbid cardiological conditions are detected and referred immediately for a proper follow up treatment. The health worker *including* the medical practitioners need to be sufficiently trained and equipped to identify abnormalities using HS. In disease like stroke and acute coronary syndrome there is a short “window period” and proper referral during this “golden hours” can save lives. Thus technological advancements in analysis of HS is called for to address, specially the rural and semi-urban India which is suffering of a huge healthcare divide.

## Acknowledgment

The authors acknowledges support of Kausik Basak, Hrushikesh Garud and the medical practitioners in-volved in the survey. SM also acknowledges the support of Texas Instruments India for supporting the research in parts.

## References

[1] S. Levine and W. Harvey, Clinical auscultation of the heart. Saunders, 1959.

[2] S. Mandal, K. Basak, K. Mandana, A. Ray, J. Chatterjee, and M. Mahadevappa, “Development of cardiac prescreening device for rural population using ultralow-power embedded system,” Biomedical Engineering, IEEE Transactions on, vol. 58, no. 3, pp. 745 –749, 2011.

[3] M. E. Tavel, “Cardiac auscultation: A glorious past – but does it have a future?” Circulation, vol. 93, no. 6, pp. 1250–1253, 1996.

[4] S. Mangione, L. Z. Nieman, E. Gracely, and D. Kaye, “The teaching and practice of cardiac auscultation during internal medicine and cardiology training. A nationwide survey,” Ann. Intern. Med., vol. 119, pp. 47–54, Jul 1993.

[5] R. Watrous, “Computer-aided auscultation of the heart: From anatomy and physiology to diagnostic decision support,” in Engineering in Medicine and Biology Society, 2006. EMBS’06. 28th Annual International Conference of the IEEE, 30 2006-sept. 3 2006, pp. 140 –143.

[6] R. L. Watrous, W. R. Thompson, and S. J. Ackerman, “The impact of computer-assisted auscultation on physician referrals of asymptomatic patients with heart murmurs,” Clinical Cardiology, vol. 31, no. 2, pp. 79–83, 2008.

[7] S. Mandal, J. Chatterjee, and A. Ray, “A new framework for wavelet based analysis of acoustical cardiac signals,” in Biomedical Engineering and Sciences (IECBES), 2010 IEEE EMBS Conference on, 30 2010-dec. 2 2010, pp. 494 –498.

[8] M. E. Tavel, “Cardiac auscultation:a glorious past–and it does have a future!” Circulation, vol. 113, no. 9, pp. 1255–1259, 2006.

[9] A. Leatham, C. Bull, and M. Braimbridge, Lecture notes on cardiology, ser. Lecture Notes Series. Blackwell Scientific, 1991.

[10] J. Hall, Guyton and Hall Textbook of Medical Physiology: With STUDENT CONSULT Online Access, ser. Guyton Physiology Series. Elsevier - Health Sciences Division, 2010.

[11] A. Bloch, J. Crittin, and A. Jaussi, “Should functional cardiac murmurs be diagnosed by auscultation or by Doppler echocardiography?” Clin Cardiol, vol. 24, pp. 767–769, Dec 2001.

[12] N. Reed, Y. Nie, and C. Mahnke, “A portable graphical representation tool for phonocardiograms,” in Engineering in Medicine and Biology Society, 2009. EMBC 2009. Annual International Conference of the IEEE, sept. 2009, pp. 3111–3114.

[13] T. R. Reed, N. E. Reed, and P. Fritzson, “Heart sound analysis for symptom detection and computer-aided diagnosis,” Simulation Modelling Practice and Theory, vol. 12, no. 2, pp. 129 –146, 2004, advances in modelling and simulation in biology and medicine.

[14] Z. Syed, D. Leeds, D. Curtis, F. Nesta, R. Levine, and J. Guttag, “A framework for the analysis of acoustical cardiac signals,” Biomedical Engineering, IEEE Transactions on, vol. 54, no. 4, pp. 651 –662, april 2007.

[15] K. Basak, S. Mandal, M. Manjunatha, J. Chatterjee, and A. Ray, “Phonocardiogram signal analysis using adaptive line enhancer methods on mixed signal processor,” in Signal Processing and Communications (SPCOM), 2010 International Conference on, july 2010, pp. 1 –5.

[16] F. A. Gari D. Clifford and P. E. M. editors., Advanced Methods and Tools for ECG Data Analysis. Artech House, 2006.

[17] Y. Akay, M. Akay, W. Welkowitz, J. Semmlow, and J. Kostis, “Noninvasive acoustical detection of coronary artery disease: a comparative study of signal processing methods,” Biomedical Engineering, IEEE Transactions on, vol. 40, no. 6, pp. 571 –578, june 1993.

[18] R. O. Duda, P. E. Hart, and D. G. Stork, Pattern classification. John Wiley & Sons, 2012.

[19] C. M. Bishop et al., Pattern recognition and machine learning. springer New York, 2006, vol. 1.

[20] A. Hyvarinen, J. Karhunen, and E. Oja, Independent component analysis. John Wiley & Sons, 2004, vol. 46.

[21] A. Hyvärinen and E. Oja, “Independent component analysis: algorithms and applications,” Neural networks, vol. 13, no. 4, pp. 411–430, 2000.

[22] A. Hyvärinen, “Fast and robust fixed-point algorithms for independent component analysis,” Neural Networks, IEEE Transactions on, vol. 10, no. 3, pp. 626–634, 1999.

[23] R. J. Martis, U. R. Acharya, H. Prasad, C. K. Chua, C. M. Lim, and J. S. Suri, “Application of higher order statistics for atrial arrhythmia classification,” Biomedical Signal Processing and Control, vol. 8, no. 6, pp. 888–900, 2013.

[24] H. Prasad, R. J. Martis, U. R. Acharya, L. C. Min, and J. S. Suri, “Application of higher order spectra for accurate delineation of atrial arrhythmia,” in Engineering in Medicine and Biology Society (EMBC), 2013 35th Annual International Conference of the IEEE. IEEE, 2013, pp. 57–60.

[25] R. J. Martis, U. R. Acharya, and H. Adeli, “Current methods in electrocardiogram characterization,” Computers in biology and medicine, vol. 48, pp. 133–149, 2014.

[26] N. Cristianini and J. Shawe-Taylor, An introduction to support vector machines and other kernel-based learning methods. Cambridge university press, 2000.

[27] B. Schölkopf and A. J. Smola, Learning with kernels: Support vector machines, regularization, optimization, and beyond. MIT press, 2002.

[28] C. Shub, “Echocardiography or auscultation? How to evaluate systolic murmurs,” Can Fam Physician, vol. 49, pp. 163–167, Feb 2003.

